# Biophysical Modeling Uncovers Transcription Factor and Nucleosome Binding on Single DNA Molecules

**DOI:** 10.1101/2025.05.13.653852

**Authors:** Lasha Dalakishvili, Husain Managori, Anaïs Bardet, Věra Slaninová, Edouard Bertrand, Nacho Molina

**Affiliations:** IGBMC | CNRS | INSERM | University of Strasbourg | 1 Rue Laurent Fries, 67404 Illkirch, France; IGH | CNRS | University of Montpellier | 141 Rue de la Cardonille, Montpellier, France

## Abstract

Gene regulation in eukaryotes emerges from a dynamic interplay between transcription factors (TFs), nucleosomes, and RNA Polymerase II (Pol II), whose competitive and cooperative binding shapes DNA accessibility and transcriptional output. Single-molecule footprinting (SMF) and long-read chromatin accessibility assays such as Fiber-seq now capture these interactions at nucleotide resolution on individual DNA molecules. However, existing computational tools remain insufficient to decode complex binding events from sparse methylation data. Here, we introduce HiddenFoot, a probabilistic modeling framework based on statistical mechanics that quantitatively infers TF, nucleosome, and Pol II occupancy profiles on single DNA molecules by systematically evaluating all thermodynamically plausible binding configurations. Applying HiddenFoot to SMF and Fiber-seq data from mouse, *Drosophila*, and human cells, we recovered known TF footprints, precisely resolved Pol II pausing, and identified extensive heterogeneity in nucleosome positioning driven by TF binding. HiddenFoot further distinguishes direct TF–TF cooperativity from nucleosome-mediated co-dependency by estimating pairwise interaction energies and comparing to null models under equilibrium. By integrating biophysical modeling with high-resolution single-molecule data, HiddenFoot offers a general, interpretable framework for dissecting regulatory logic in native chromatin with base-pair precision.

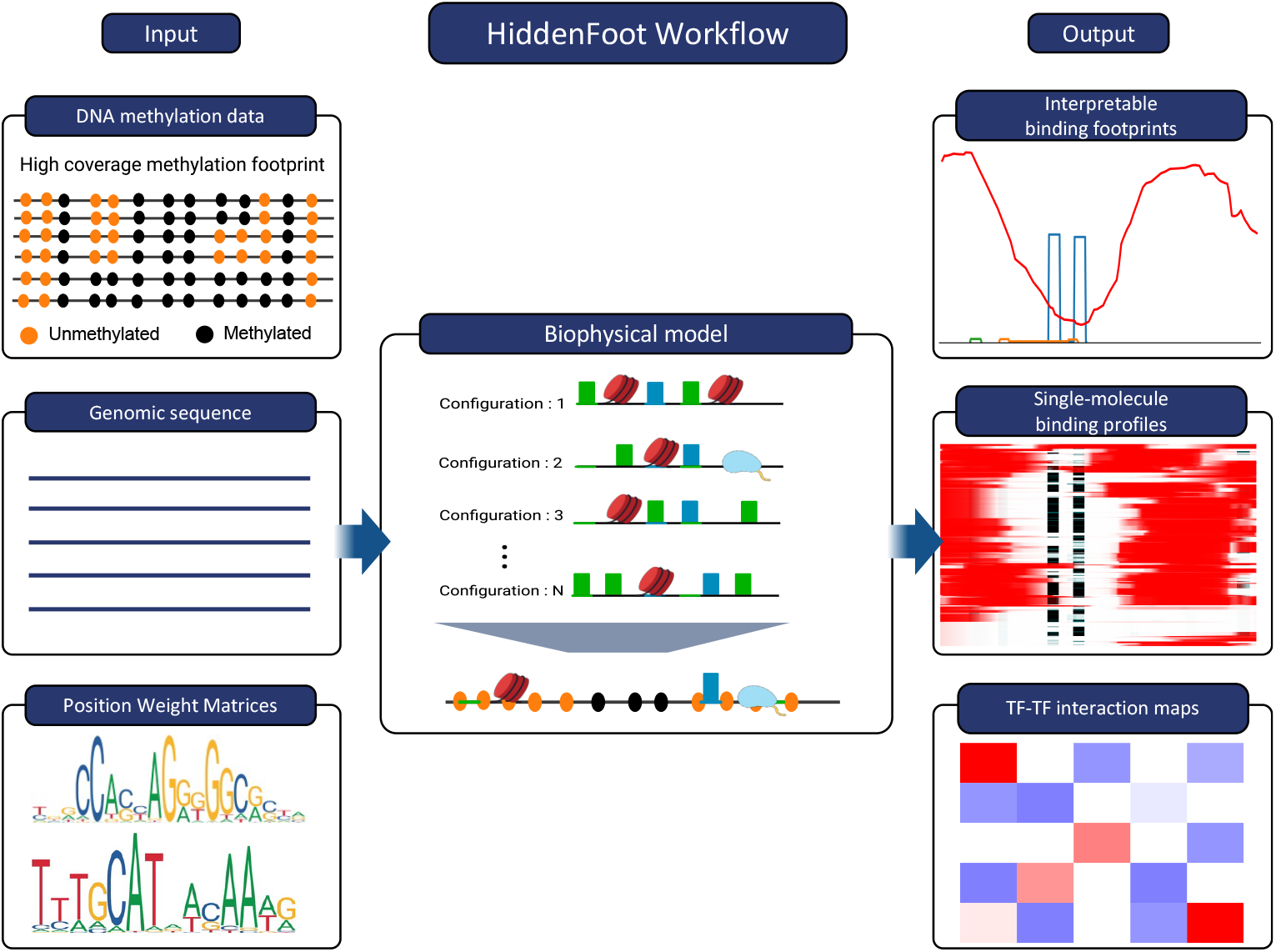

## Introduction

In eukaryotic genomes, the genetic material is compacted into chromatin, a dynamic structure in which DNA is wrapped around histone octamers to form nucleosomes [1]. This packaging serves as both a physical barrier and a regulatory scaffold for gene expression. Transcription factors (TFs) must gain access to their target sequences within chromatin to initiate transcription, often in the context of a highly competitive and cooperative molecular environment [2]. TFs frequently function in concert, displacing or repositioning nucleosomes to facilitate binding at cis-regulatory elements such as promoters and enhancers [3, 4]. At promoters, this interplay extends to RNA Polymerase II (Pol II), whose recruitment and pausing are tightly coordinated with local chromatin architecture [5, 6]. While it is well established that TF binding and nucleosome positioning are interdependent processes, the underlying mechanisms by which they coordinate and compete for DNA remain incompletely understood.

Conventional techniques to study protein-DNA interactions, such as chromatin immunoprecipitation followed by sequencing (ChIP-seq), provide genome-wide snapshots of TF or Pol II occupancy [5, 7]. However, these methods are inherently limited by their reliance on bulk cell populations. As a result, they average over diverse molecular states, masking important regulatory heterogeneity and obscuring the transient or combinatorial nature of binding events. For example, ChIP-seq does not resolve whether two TFs bind simultaneously to the same DNA molecule or whether they act in mutually exclusive fashion. Similarly, MNase-seq and ATAC-seq measure nucleosome positioning and chromatin accessibility across populations but cannot resolve the variability of these features at the single-molecule level [8, 9].

To overcome these limitations, single-molecule chromatin accessibility mapping technologies such as Single-Molecule Footprinting (SMF) [10] and single-molecule chromatin fiber sequencing (Fiber-seq) [11] have emerged as powerful approaches to profile chromatin architecture at nucleotide resolution on individual DNA molecules. These techniques use exogenous methyltransferases to label accessible cytosines (SMF) or adenines (Fiber-seq), enabling detection of TF and nucleosome footprints without DNA fragmentation. By avoiding cell averaging, SMF reveals molecular heterogeneity in TF occupancy, cooperative binding, nucleosome phasing, and transcriptional engagement across the genome [12–16]. However, extracting biologically meaningful protein-DNA interactions from SMF methylation patterns remains a major computational challenge. Particularly in regulatory regions where many TF motifs are densely packed and nucleosome positioning is variable, inferring the specific binding configurations that produced a given methylation pattern is a nontrivial inverse problem. Thus, there is a need for computational methods capable of analyzing SMF-like data in a mechanistic and interpretable manner.

To address this, we present HiddenFoot, a probabilistic modeling framework that integrates single-molecule methylation data with a biophysical model of protein-DNA binding. Inspired by Boltzmann-Gibbs statistical mechanics, HiddenFoot computes the probability of all sterically allowed configurations of TFs and nucleosomes on a given DNA sequence, accounting for sequence-specific binding affinities via Position Weight Matrices (PWMs), local protein concentrations, and molecular competition for DNA. By marginalizing over configurations and conditioning on observed methylation data, HiddenFoot infers the posterior probability of the binding state of each base pair at single-molecule resolution. This approach allows us to distinguish bound and unbound states for individual TFs and nucleosomes, resolve co-binding and exclusion patterns, and quantify dependencies between TF pairs.

In contrast to statistical HMM-based models or heuristic clustering approaches [14, 17], HiddenFoot is grounded in thermodynamic principles and explicitly models protein-DNA competition. This enables mechanistically interpretable inference of occupancy states and interaction energies at single-molecule resolution. It provides base-pair–level estimates of occupancy for hundreds of proteins across thousands of molecules, and its statistical formulation enables both global parameter estimation and per-molecule posterior inference.

We applied HiddenFoot to three distinct datasets to demonstrate its generalizability and interpretability across biological systems and assay platforms. In mouse embryonic stem cells (mESCs), HiddenFoot distinguished direct TF–TF cooperativity from nucleosome-mediated co-dependencies, offering mechanistic insights into regulatory interactions at base-pair resolution. In engineered HeLa cells containing an HIV-1 reporter, HiddenFoot resolved Pol II occupancy on individual DNA molecules and revealed chromatin reorganization in response to transcriptional inhibition. Finally, in *Drosophila* S2 cells, analysis of Fiber-seq data uncovered widespread nucleosome positioning heterogeneity and showed how TF binding locally shapes chromatin structure. Together, these results highlight HiddenFoot’s capacity to extract fine-grained regulatory information from sparse single-molecule measurements across diverse biological and experimental contexts.

In summary, HiddenFoot unlocks the full potential of single-molecule footprinting by offering a unified, physics-informed approach to reconstruct the combinatorial logic of TF, nucleosome and Pol II binding. While initially developed for methylation-based SMF and Fiber-seq data, HiddenFoot’s thermodynamic framework is, in principle, easily adaptable to any other single-molecule chromatin accessibility mapping approach based on non-disruptive chemical tagging. Finally, by moving beyond population averages to model molecular configurations on individual DNA templates, HiddenFoot has the potential to shed new light on the regulatory grammar of chromatin and offering a foundation for decoding the molecular basis of gene regulation with unprecedented resolution.

## Results

### An Efficient and Interpretable Biophysical Approach for SMF Analysis

To resolve TF and nucleosome binding on individual DNA molecules from sparse methylation data, we developed HiddenFoot, a biophysical modeling framework that integrates DNA sequence specificity, local protein concentrations, and SMF measurements. Rather than relying on heuristic filters or local signal patterns, HiddenFoot explicitly models the competition between proteins for DNA binding under thermodynamic equilibrium, enabling accurate inference of protein occupancy at single-molecule resolution.

At the core of HiddenFoot is a statistical mechanical model that evaluates all possible non-overlapping configurations *C* of bound proteins (TFs and nucleosomes) along a given DNA sequence *S* (see Fig. 1A). The probability of each configuration is determined using the Boltzmann-Gibbs distribution [18, 19]:

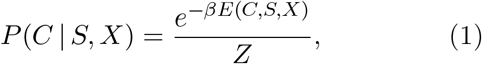

where *β* is the inverse thermal energy, *X* denotes the set of local protein concentrations, and *E*(*C, S, T*) is the total energy of the configuration. This total energy is computed as the sum of the individual binding energies of all proteins in the configuration *C*, with each energy term derived from PWMs. The protein concentrations determine the effective chemical potentials, which modulate the statistical weight of each protein’s binding to the overall configuration likelihood. These concentrations are tunable parameters learned from the data, and they reflect the relative contribution of each protein to explaining the observed methylation patterns (see Methods). The normalization constant *Z* is the partition function summing over all thermodynamically allowed (i.e., non-overlapping) binding configurations.

**Figure 1:**
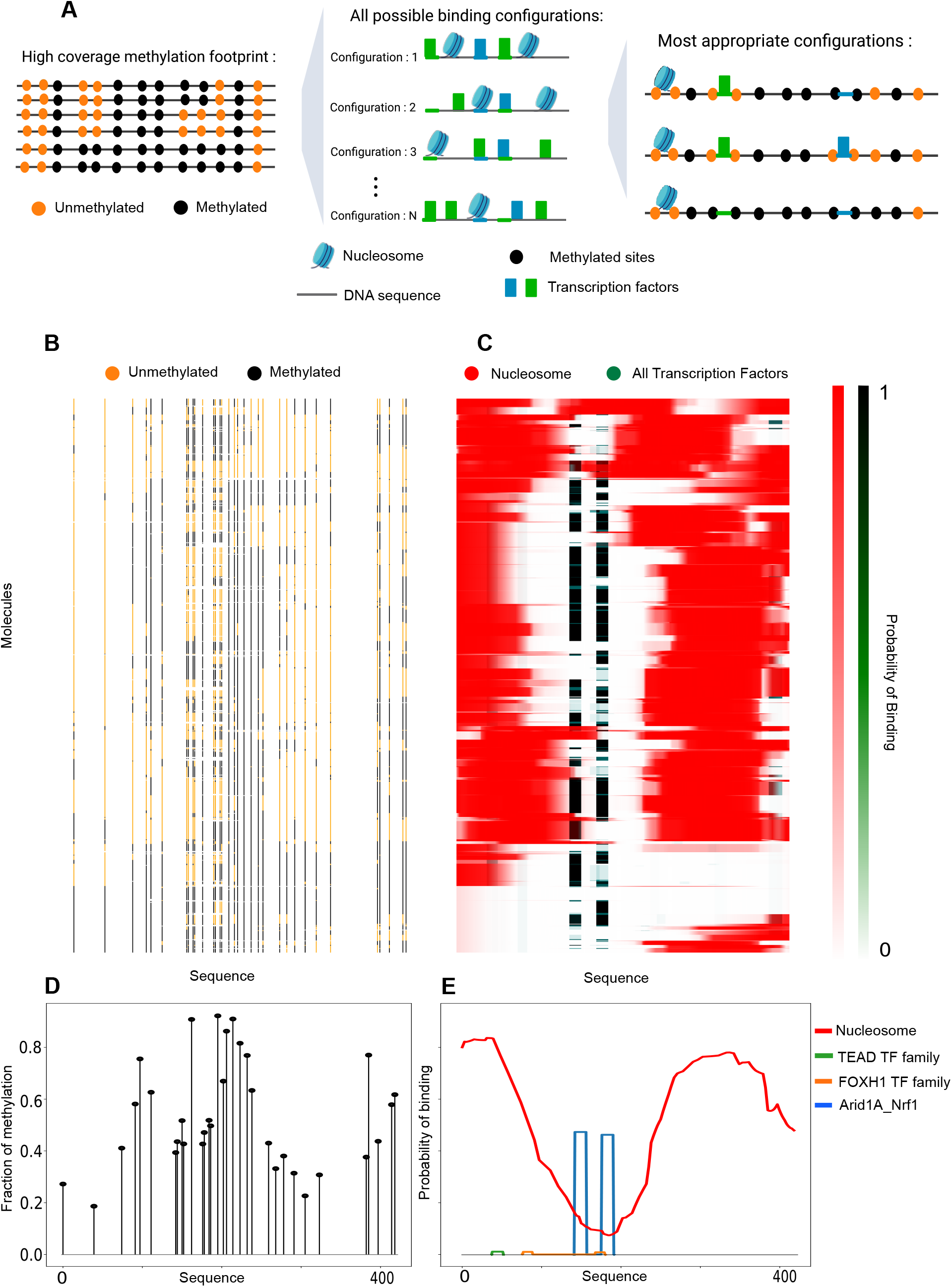
HiddenFoot infers TF and nucleosome binding profiles from single-molecule DNA methylation data. (A) Schematic overview of the HiddenFoot framework. Single-molecule methylation patterns are interpreted to evaluate all possible binding configurations of TFs and nucleosomes, identifying the most likely occupancy states. (B) Raw methylation data across all molecules for a representative amplicon region. Orange indicates unmethylated (protein-bound) sites, and black indicates methylated (accessible) sites. (C) Clustered binding probability profiles inferred by HiddenFoot, showing nucleosome occupancy in red and combined TF occupancy in green across molecules. (D) Average methylation profile across the region, aggregated over all molecules. (E) Average binding profiles inferred by HiddenFoot, showing nucleosome occupancy (red) and individual TFs in distinct colors.

To connect these biophysical binding configurations to experimentally observed SMF methylation patterns *M*, HiddenFoot defines a likelihood function *P* (*M* |*C, Q*), which describes the probability of observing a methylation pattern *M* given a binding configuration *C* and a pair of occupancy-dependent metyhlation parameters *Q* = {*q*_*U*_, *q*_*B*_}. Here, *q*_*U*_ is the probability that a site is methylated when unbound, and *q*_*B*_ is the probability that a site is methylated when bound. These parameters account for the stochasticity of the methylation process and the imperfect correspondence between protein occupancy and methylation state.

The resulting likelihood of observing a methylation pattern *M* for a given DNA sequence *S*, protein concentration set *T*, and methylation parameters *Q*, marginalizing over all possible configurations *C*, is:

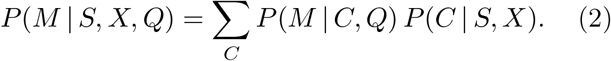

Critically, the sum over exponentially many non-overlapping configurations can be efficiently computed using dynamic programming (see Methods), allowing the model to scale to long DNA regions with hundreds of potential binding sites.

Model parameters, including the protein concentrations and methylation probabilities, are estimated by maximizing the likelihood of the observed data using stochastic gradient descent (SGD). Owing to the dynamic programming formulation, analytical gradients can be computed efficiently, enabling tractable optimization even across thousands of molecules. In addition, HiddenFoot estimates posterior uncertainties on the parameters by sampling from the posterior distribution using a Metropolis-Hastings Markov Chain Monte Carlo (MCMC) algorithm (see Methods).

Finally, once model parameters are fitted, HiddenFoot computes the marginal posterior probability that each base is bound by a specific protein using an efficient forward–backward algorithm (see Methods). This enables high-resolution inference of occupancy landscapes across individual DNA molecules by leveraging the same dynamic programming framework used for computing the partition function and the mariginal likelihood.

### HiddenFoot infers TF and nucleosome binding profiles at single-molecule DNA resolution

We evaluated HiddenFoot on SMF data from mouse embryonic stem cells (mESCs), consisting of 246 high-coverage amplicons targeting known regulatory regions [13]. These regions span a diversity of chromatin contexts and TF motifs, providing a rigorous benchmark for assessing model performance. HiddenFoot was run independently on each amplicon using 258 clustered PWMs from the JASPAR database [20], allowing region-specific inference of protein concentrations and binding configurations, and enabling parallelized analysis across the dataset.

To illustrate the interpretability and resolution of the inferred binding profiles, we show results for one representative amplicon region in Fig. 1. The raw methy-lation data (Fig. 1B) display binary patterns across individual DNA molecules, where black marks indicate methylated (accessible) cytosines and orange marks indicate unmethylated (protected) cytosines, presumably bound by proteins. The average methylation profile across all molecules in the region (Fig. 1D) reveals a characteristic landscape of protected and accessible sites, consistent with phased nucleosomes and focal TF binding.

However, these aggregate signals alone are insufficient to disentangle the contributions of specific TFs and nucleosomes. HiddenFoot leverages the underlying sequence information and thermodynamic modeling to decode these patterns. Fig. 1E shows the average binding profiles inferred by the model, which separate nucleosome occupancy (red) from individual TF footprints (colored lines). The model accurately recovers nucleosome-depleted regions harboring discrete TF binding sites and phased nucleosome arrays flanking them, hallmarks of active regulatory elements.

Importantly, HiddenFoot goes beyond population averages to resolve protein-DNA interactions on a per-molecule basis. Fig. 1C displays a heatmap of the inferred binding probabilities for each individual molecule in the amplicon, clustered by similarity. The model distinguishes nucleosome binding (red) from combined TF binding (green), revealing heterogeneity in binding patterns across molecules. This level of interpretability is unique: to our knowledge, HiddenFoot is the first approach capable of transforming sparse, noisy SMF data into mechanistically meaningful binding maps at single-molecule resolution.

To assess the overall performance of the model in capturing methylation landscapes, we compared the average methylation levels predicted by HiddenFoot to the observed experimental values across all cytosines in the 246 amplicon regions. As shown in Supplementary Fig. S1A, the predicted and observed methylation levels exhibit a strong linear correlation at the single-base level, indicating that the model faithfully reconstructs global accessibility patterns from its inferred binding states. This agreement holds consistently across individual amplicons, with Pearson correlation coefficients exceeding 0.8 for the majority of regions (Supplementary Fig. S1B). The high reconstruction accuracy demonstrates that HiddenFoot can quantitatively recapitulate the observed methylation signal as a result of TF and nucleosome binding.

Beyond reconstruction accuracy, we also evaluated the robustness of parameter inference using posterior sampling. For each amplicon, we estimated the posterior distributions of key model parameters (including TF and nucleosome concentrations as well as the metylation probabilities) using Metropolis-Hastings MCMC. Supplementary Fig. S2 summarizes the posterior variability across all amplicons using the coefficient of variation (CV), defined as the standard deviation divided by the mean. Nucleosome concentration estimates are particularly well-constrained, with CVs typically below 10%, while TF concentrations and *β* show moderate variability, with most CVs falling below 30%. These results indicate that the model is not only expressive enough to fit diverse methylation patterns but also identifiable, providing confidence in the biological interpretability of the inferred concentrations and metyhlation probabilities.

In summary, by integrating statistical mechanics, sequence specificity, and methylation state likelihoods, HiddenFoot enables quantitative dissection of chromatin occupancy from SMF data. Unlike traditional accessibility-based footprinting, which lacks resolution and mechanistic grounding, HiddenFoot provides a principled framework to resolve competitive binding events molecule-by-molecule.

### HiddenFoot Classifies Bound Molecules and Recovers Biologically Meaningful TF Footprints

To evaluate the biological validity of HiddenFoot’s TF binding predictions, we applied the model to a genome-scale SMF dataset generated by targeted DNA capture in mESCs. This dataset encompasses over 297,000 genomic regions, covering approximately 2% of the genome, and includes a large fraction of candidate cis-regulatory elements (CREs) that are accessible in mESCs [13]. The comprehensive coverage and depth of this dataset allowed us to rigorously benchmark HiddenFoot’s predictive power against orthogonal experimental data.

Our validation strategy focused on two key objectives: (i) determining whether HiddenFoot-derived binding probabilities can stratify individual molecules into biologically meaningful bound and unbound classes, and (ii) assessing the correlation between predicted binding probabilities and ChIP-seq enrichment at TF-specific binding sites. To this end, we first selected molecules overlapping known TF motifs and stratified them either by ChIP-seq signal rank (using only molecules that mapped to the top 10% of ChIP-seq-enriched regions) or by HiddenFoot’s inferred per-molecule binding state. As shown in Figure 2A–D, averaging the methylation profiles of molecules within top-ranked ChIP-seq regions yields footprints of varying clarity: while CTCF and REST show detectable protection patterns, NRF1 and ESRRB do not produce clear signatures. In contrast, when molecules are stratified based on HiddenFoot-predicted occupancy (see Methods), robust footprints emerge for all TFs (Fig. 2E–H), including NRF1 and ESRRB. This improvement stems from HiddenFoot’s ability to disambiguate bound versus unbound molecules within the same region, highlighting a major limitation of standard signal-based aggregation approaches that pool both bound and unbound molecules indiscriminately.

**Figure 2:**
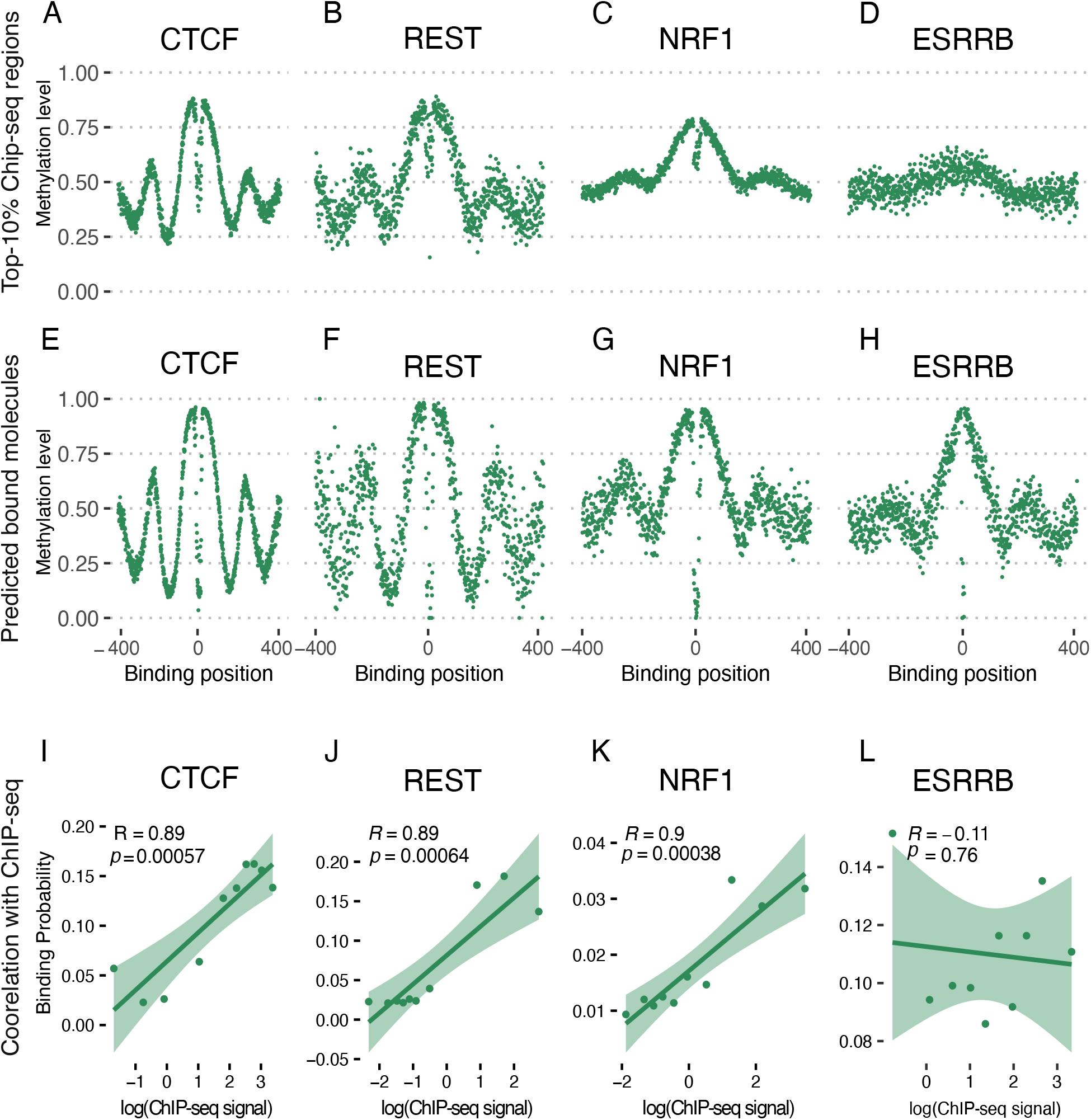
HiddenFoot detects methylation footprints in mouse embryonic stem cells (mESC). **(A–D)**: Average methylation level at the binding site of the top 10% ChIP-seq enriched molecules for CTCF (A), REST (B), NRF1 (C), and ESRRB (D). Each dot represents the average methylation level at a given position relative to the TF motif center. **(E–H)**: Average methylation levels at the same TF motifs, stratified using HiddenFoot predictions. Molecules are divided based on inferred binding state. Clear footprint-like patterns are visible, particularly for CTCF and REST. **(I–L)**: Pearson correlation between the average HiddenFoot-predicted binding probabilities and log-transformed ChIP-seq signal, binned into 10 groups. Each dot corresponds to one bin. CTCF (I), REST (J), and NRF1 (K) show high correlation (*R >* 0.85), while ESRRB (L) does not.

More importantly, HiddenFoot binding probabilities correlate strongly with independent ChIP-seq datasets. For each motif-centered region, we computed the average TF binding probability and compared it to the log-transformed ChIP-seq signal across binned quantiles. The resulting Pearson correlation coefficients exceeded 0.85 for CTCF, REST, and NRF1 (Fig. 2I– K), validating the quantitative concordance between model-based predictions and ChIP-seq occupancy. Interestingly, ESRRB exhibited no detectable correlation (Fig. 2L), despite showing a weak footprint signal. This discrepancy may reflect technical limitations of the ESRRB ChIP-seq dataset, which has a low signal-to-noise ratio and limited read depth, or it could indicate rapid TF turnover at ESRRB sites, leading to weak or transient protection signals in SMF data. This observation illustrates both a potential limitation of the footprinting approach (its sensitivity to TF residence time) and the value of integrating multiple orthogonal datasets for robust occupancy assessment.

In addition to validating TF-specific binding predictions, we tested whether HiddenFoot’s inferred DNA accessibility probabilities correlate with genome-wide chromatin accessibility measured by MNase-seq and ATAC-seq. As shown in Supplementary Fig. S3, HiddenFoot predictions of accessibility exhibit strong concordance with both assays, with Pearson correlation coefficients of *R* = 0.97 and *R* = 0.96, respectively. These results underscore the model’s capacity to recapitulate key chromatin features from sparse, single-molecule methylation data, further reinforcing the biological interpretability of its predictions.

Together, these analyses demonstrate that HiddenFoot not only improves TF footprint detection through molecule-level classification, but also provides quantitative predictions that align with ChIP-seq and chromatin accessibility data. These findings establish HiddenFoot as a robust and generalizable tool for decoding TF and nuclosome binding landscapes from single-molecule measurements.

### HiddenFoot Distinguishes Nucleosome-Mediated and Direct Physical TF Cooperativity

The single-molecule resolution of HiddenFoot enables systematic quantification of TF co-occupancy and interaction patterns across genomic loci. To distinguish direct physical TF-TF cooperativity from indirect co-binding effects driven by nucleosome exclusion, we inferred an interaction energy parameter *E*_12_ for each pair of TFs. This parameter captures whether TFs bind independently, repulsively, or cooperatively on the same DNA molecule.

The interaction energy *E*_12_ is estimated from HiddenFoot-inferred marginal binding probabilities across all molecules using a probabilistic occupancy model (see Methods). For each pair of TF binding sites, we compute the expected number of molecules in each co-binding state—both unbound (*N*_00_), only TF1 bound (*N*_10_), only TF2 bound (*N*_01_), and both bound (*N*_11_)—and fit a statistical mechanics model that assigns likelihoods to each configuration based on individual binding energies *E*_1_, *E*_2_, and the interaction term *E*_12_. Positive values of *E*_12_ indicate cooperative binding, negative values suggest mutual exclusion, and values near zero correspond to independent behavior.

Fig. 3A–C illustrates representative patterns of TF-TF co-dependency: correlated binding (both TFs tend to bind together), anticorrelated binding (one binds when the other does not), and independent binding. These patterns are evident in real data from mESCs (Fig. 3D–F). For example, Arid1A_Nrf1 and Gmeb1 display frequent co-binding (correlated), ZNF627 and ZNF611 are largely mutually exclusive, likely due to nucleosome competition; while ZNF594 and ZNF77 bind independently across molecules.

**Figure 3:**
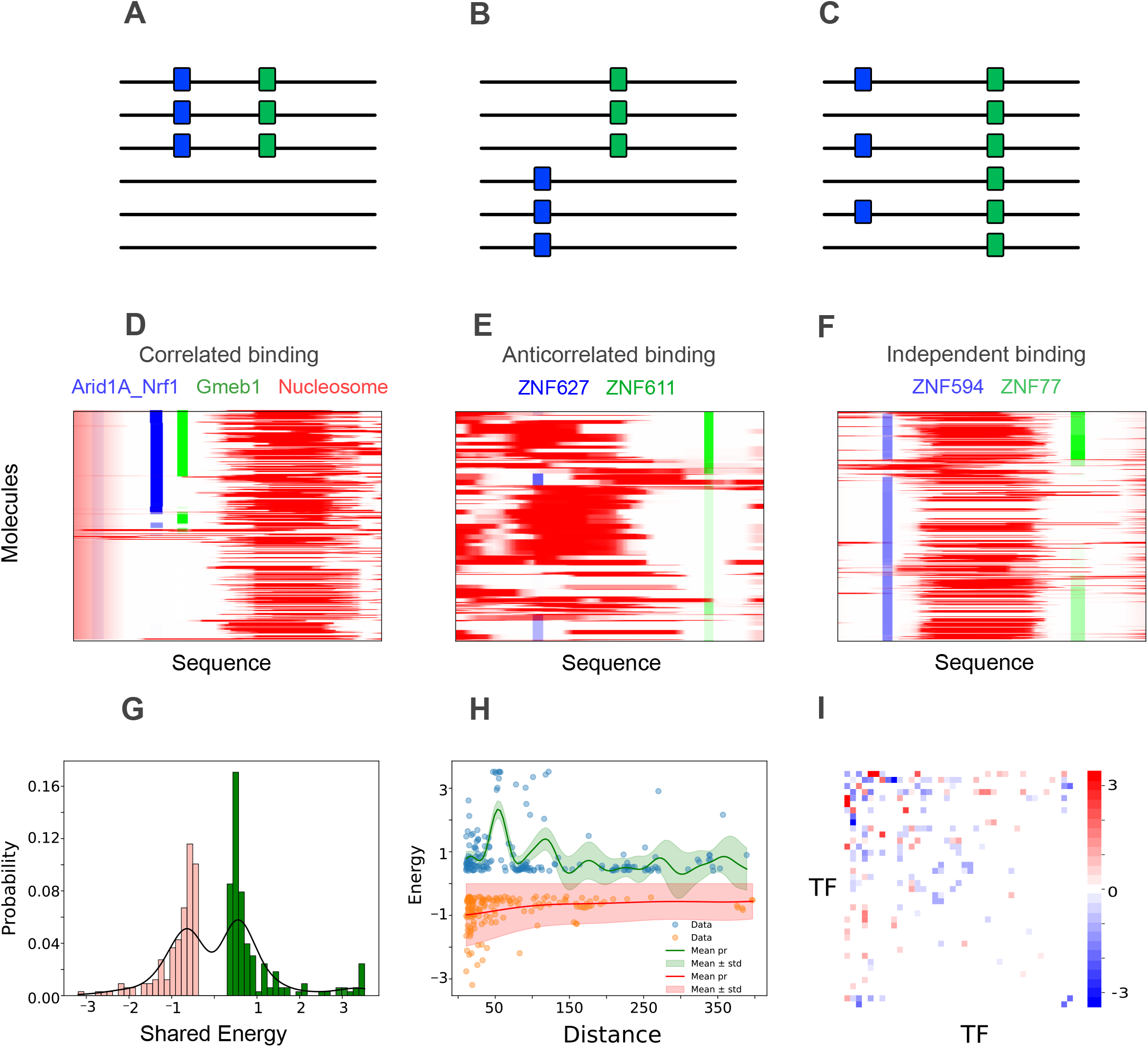
HiddenFoot infers and differentiates nucleosome-mediated and physical TF-TF cooperativity. **(A–C)**: Schematic examples of TF-TF co-dependency patterns: correlated (A), anticorrelated (B), and independent (C) binding. **(D–F)**: Real data examples from mESCs. (D) Arid1A_Nrf1 and Gmeb1 show co-binding on the same molecules (correlated). (E) ZNF627 and ZNF611 are anticorrelated, but ZNF611 binding regions are occupied by nucleosomes, suggesting indirect exclusion. (F) ZNF594 and ZNF77 bind independently. **(G)**: Distribution of inferred TF-TF shared energies. Positive values reflect cooperative interactions; negative values suggest mutual exclusion. **(H)**: TF-TF interaction energy as a function of distance. Peaks near 10, 50, and 150 bp support structural chromatin influence on cooperation. **(I)**: Heatmap comparing shared energies between TF pairs in real vs. simulated data. TF pairs with divergent values likely involve direct physical interactions.

The genome-wide distribution of inferred *E*_12_ values reveals that most TF pairs cluster around zero, consistent with independent binding, while a notable fraction exhibits substantial positive or negative interaction energies (Fig. 3G). The spatial distribution of inferred co-binding may reflect chromatin structure: whereas negative interaction energies decay smoothly with increasing distance between TF binding sites, positive energies display characteristic peaks at ~10 bp, ~ 50 bp, and ~140 bp (Fig. 3H). These distances may correspond to the DNA helical pitch, phased TF spacing near nucleosome-depleted regions, and the typical footprint of a nucleosome.

To disentangle direct TF–TF cooperativity from nucleosome-mediated interactions, we simulated binding profile datasets using the binding parameters inferred by HiddenFoot, but without conditioning the binding configurations on the observed methylation data. These simulations represent equilibrium binding configurations in the absence of external remodeling forces or direct protein–protein contacts, while still accounting for steric exclusion between bound proteins. By comparing the shared energies inferred from real and simulated data, we can attribute deviations to bona fide physical cooperativity. If a TF pair exhibits similar co-binding behavior in both real and simulated datasets, the interaction is likely mediated by chromatin accessibility and steric exclusion. In contrast, if a stronger co-dependency is observed in the real data compared to the simulated data, it likely reflects a direct molecular interaction.

This comparison is visualized in a heatmap of shared interaction energies for all TF pairs in the real versus simulated datasets (Fig. 3I). In the simulated data, TF–TF co-dependencies arise solely from nucleosome-mediated effects, since no direct interaction energies are modeled. Thus, TF pairs with similar shared energies in both real and simulated data likely reflect chromatin-driven co-occupancy. In contrast, TF pairs that show strong interaction energies in the real data but not in simulation are indicative of direct physical cooperativity. Notably, several of these predicted interactions correspond to biologically supported TF pairs (see Supplementary Table 1). For example, MEIS1 and SOX4 are known to co-occupy super-enhancers during retinoic acid–induced differentiation [21, 22], while members of the SOX and NFI families have been reported to interact in large-scale protein–protein interaction studies [23]. Furthermore, FOXH and OTX family transcription factors co-regulate common genes by binding to shared regulatory elements and activating lineage-specific gene expression during early development [24]. Finally, self-interactions involving PITX3, OTX2, MEIS1, and NFIX are also supported by evidence of cooperative binding in transcriptional complexes [25–28]. Together, these observations support the ability of HiddenFoot to disentangle structural influences from intrinsic biochemical TF–TF interactions and to identify biologically meaningful combinatorial regulation in native chromatin contexts.

### HiddenFoot Reveals Pol-II Footprints in HIV-1 Promoter

To assess whether HiddenFoot can accurately resolve Pol II occupancy at single-molecule resolution, we turned to the well-characterized HIV-1 promoter as a model system. The integrated HIV-1 provirus provides a unique setting to explore transcriptional regulation, as its activity is tightly controlled by the viral transactivator Tat and is highly sensitive to perturbations in the transcription initiation machinery [29]. Using available SMF data from engineered HeLa cells containing a single-copy HIV-1 reporter and constitutively expressing the viral activator Tat [30], we tested whether HiddenFoot could detect Pol II footprints on individual DNA molecules and capture dynamic changes in promoter architecture under transcriptionally active and repressed conditions.

Pol II was modeled as a non-sequence-specific factor occupying 40 base pairs of DNA, allowing it to bind competitively with nucleosomes and TFs without requiring a DNA motif. HiddenFoot was run on SMF data from cells under two conditions: active transcription (High Tat) and transcriptional inhibition using the initiation blocker Triptolide (TLD). In both cases, we inferred per-molecule occupancy profiles for Pol II, nucleosomes, and TFs across the HIV-1 promoter.

Figure 4A–B shows heatmaps of inferred occupancy for individual molecules, sorted by total nucleosome coverage. Under High Tat conditions, Pol II footprints (blue) are prominently localized just downstream of the transcription start site (TSS), coinciding with local nucleosome depletion. This pattern is consistent with active transcription initiation and aligns with prior observations of Pol II pausing at the HIV-1 promoter [31]. Interestingly, we also observe a smaller but distinct symmetric Pol II footprint located upstream, which may reflect divergent transcription initiation at the promoter [32, 33]. Notably, our analysis clearly resolves the Nuc-0 and Nuc-1 positions flanking the promoter, revealing interesting heterogeneity in their occupancy across individual molecules.

**Figure 4:**
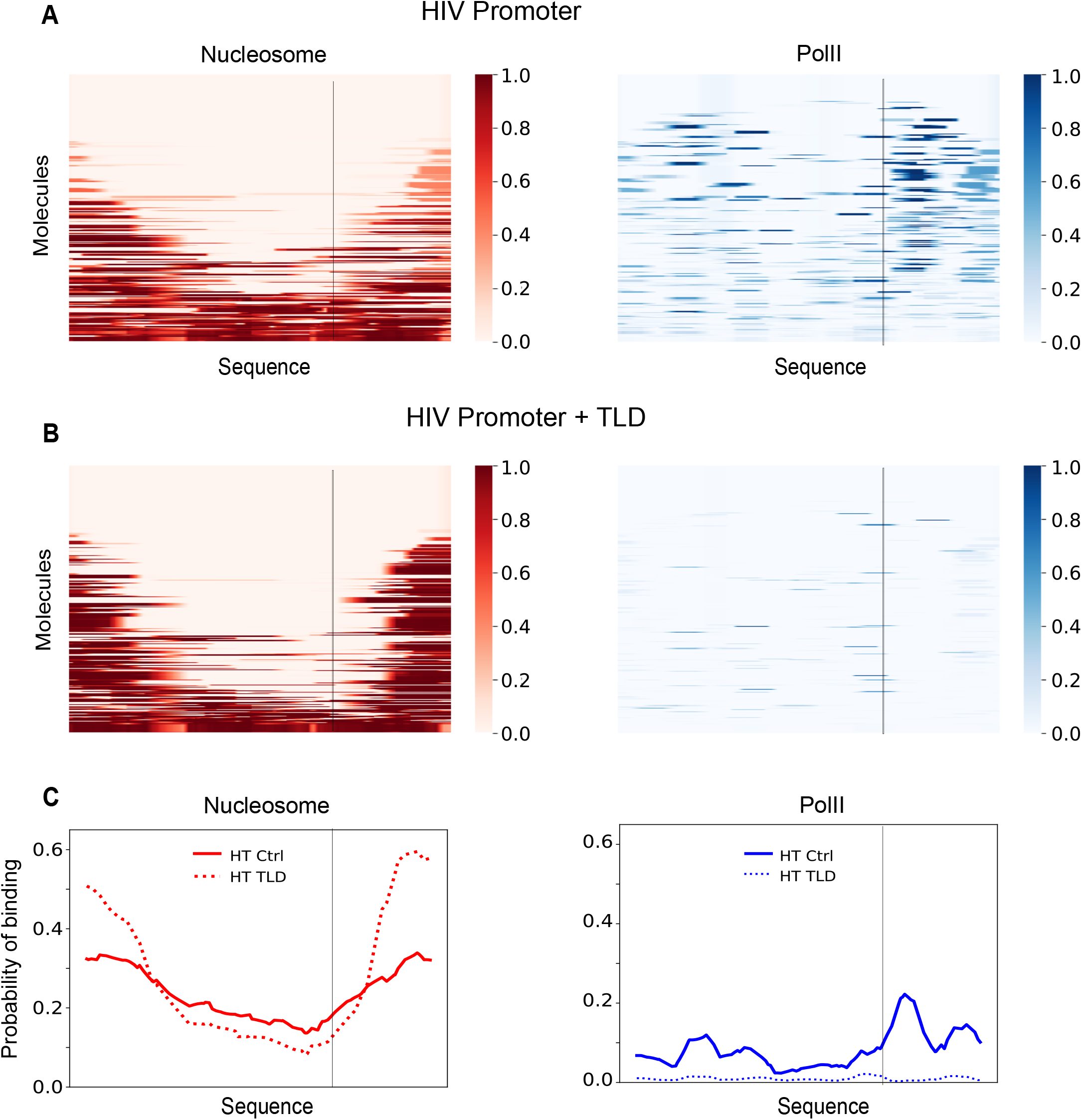
HiddenFoot resolves Pol II and nucleosome occupancy at the HIV-1 promoter under transcriptionally active and inhibited conditions. **(A–B)** Single-molecule occupancy heatmaps estimated by HiddenFoot for the HIV-1 promoter in cells with high transcriptional activity (A) and following Triptolide (TLD) treatment (B). Each row corresponds to an individual DNA molecule, sorted by total nucleosome occupancy. HiddenFoot estimates the binding probabilities of nucleosomes (red) and RNA Pol II (blue), revealing distinct chromatin configurations across molecules. Under high transcription, Pol II footprints are frequent and concentrated downstream the TSS (black line). Upon TLD treatment, Pol II occupancy is markedly reduced, while nucleosome coverage increases. **(C)** Average binding profiles across all molecules in high transcriptional activity (solid lines) and following TLD treatment (dashed lines). Pol II occupancy disappears under TLD, consistent with global inhibition of transcription initiation.

In contrast, following TLD treatment, Pol II occupancy is strongly reduced across molecules, while nucleosome occupancy increases, particularly at Nuc-0 and Nuc-1 positions. This shift indicates that transcription initiation is blocked and that nucleosomes reoccupy previously accessible regions. These effects are further confirmed by average binding profiles (Fig. 4C), where the sharp Pol II peak at the TSS observed in High Tat cells is nearly abolished in the presence of TLD.

Together, these results highlight HiddenFoot’s ability to resolve transcription initiation events, quantify Pol II occupancy at the single-molecule level, and distinguish changes in promoter architecture driven by perturbations to the transcriptional machinery.

### HiddenFoot Resolves TF-Induced Nucleosome Heterogeneity in Fiber-seq Data

To demonstrate the generality of HiddenFoot beyond SMF and its ability to interpret data from diverse experimental platforms, we applied the model to a Fiber-seq dataset generated in *Drosophila* S2 cells [11]. Fiber-seq provides long-range, single-molecule resolution of chromatin architecture by detecting DNA accessibility through exogenous adenine methylation, enabling the reconstruction of nucleosome and protein occupancy across extended genomic regions. This analysis allowed us to validate HiddenFoot’s performance on an independent assay and investigate nucleosome positioning heterogeneity influenced by TF binding.

As in the SMF analysis, we first validated HiddenFoot’s TF binding predictions against orthogonal datasets. Fig. S4 shows that HiddenFoot accurately detects methylation footprints for several TFs (CTCF, MAX, and CLAMP) when stratifying molecules by inferred binding states. Compared to naive aggregation based on ChIP-seq rank, the model-derived foot-prints are sharper and more localized, reflecting HiddenFoot’s ability to resolve true bound molecules. Furthermore, TF binding probabilities inferred by HiddenFoot strongly correlate with ChIP-seq signal intensities across genomic regions (Fig. S4G–I), with Pearson correlation coefficients of *R* = 0.88 for CTCF, *R* = 0.78 for MAX, and *R* = 0.84 for CLAMP. These results support the model’s accuracy in predicting TF occupancy from Fiber-seq methylation data. Similarly, model-inferred DNA accessibility is highly concordant with bulk chromatin accessibility profiles from MNase-seq and ATAC-seq, with correlation coefficients of *R* = 0.73 and *R* = 0.89, respectively (Fig. S5). These findings confirm that HiddenFoot captures biologically meaningful features of chromatin structure in Fiber-seq datasets.

We then leveraged HiddenFoot’s probabilistic per-molecule inference to investigate nucleosome positioning heterogeneity across thousands of individual DNA molecules. Fig. 5 illustrates three representative loci centered on MAX TF binding motifs. Each panel displays per-molecule nucleosome occupancy probabilities (in red) overlaid with MAX binding probabilities (in black). These heatmaps reveal substantial molecule-to-molecule variation in nucleosome positioning, even within the same genomic region. While some molecules exhibit phased arrays of nucleosomes flanking a nucleosome-depleted region, others show irregular or partially disorganized nucleosome configurations. This shows that HiddenFoot provides a powerful framework for studying how individual TF binding events may influence nucleosome organization at single-molecule resolution.

**Figure 5:**
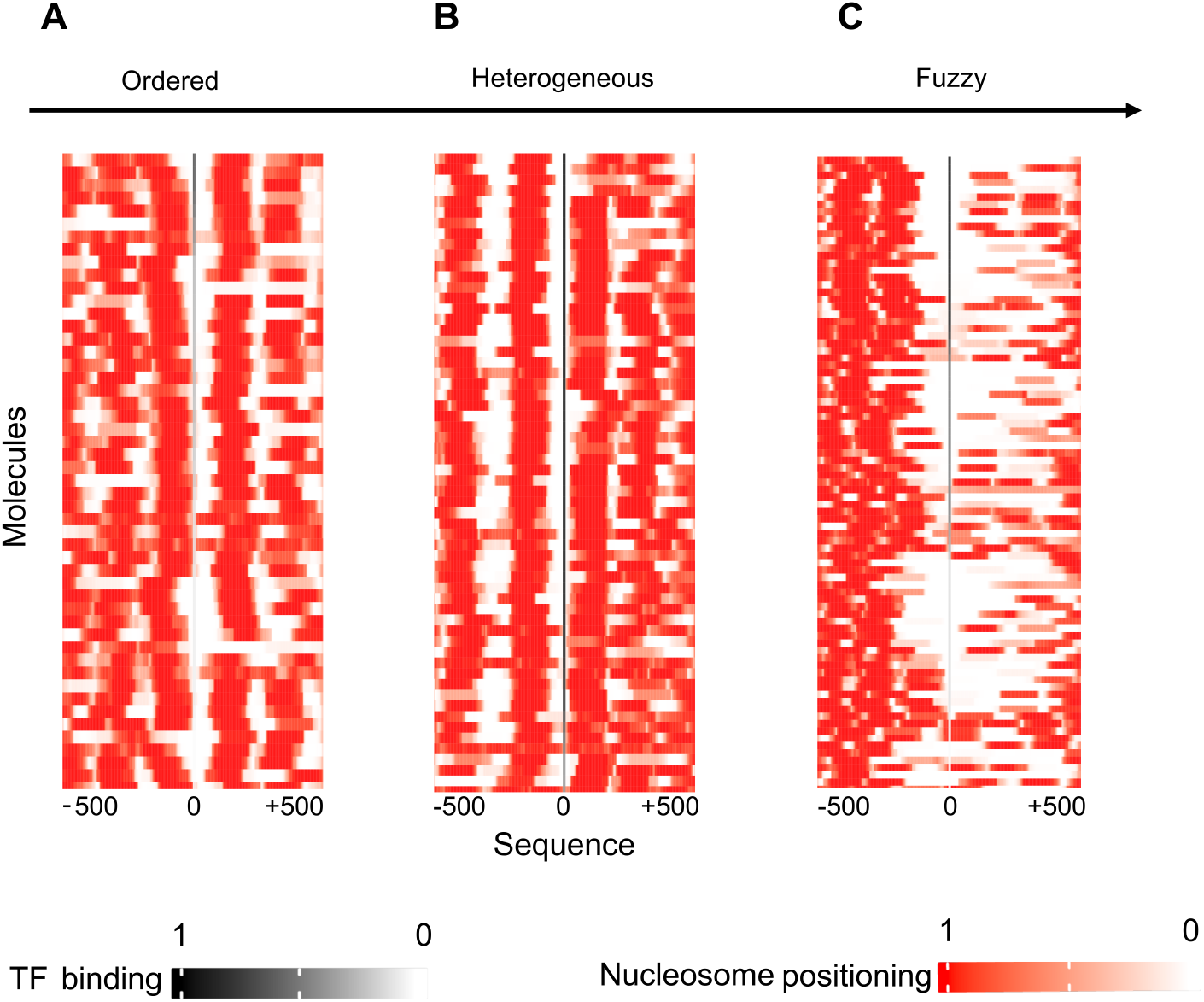
HiddenFoot reveals heterogeneity in nucleosome positioning on long DNA molecules from Fiber-seq experiments. Heatmaps show nucleosome occupancy probabilities (red) across individual long DNA molecules, centered on MAX TF motifs in three distinct genomic regions. Inferred MAX binding probabilities are overlaid in black. This visualization highlights substantial molecule-to-molecule variability in nucleosome positioning and illustrates how MAX binding locally shapes chromatin organization across different genomic contexts.

To better assess nucleosome heterogeneity associated with MAX binding globally, we clustered molecules genome-wide based on their nucleosome occupancy profiles centered on MAX motifs (Fig. S6). We stratified molecules into those predicted to be bound or unbound by MAX. The resulting heatmaps show interesting differences: molecules with bound MAX exhibit strongly phased nucleosome arrays and clear central depletion, while unbound molecules display more variable, less organized nucleosome configurations. These results indicate that MAX binding may not only perturb nucleosome positioning locally, but possible could also impose structural order over surrounding chromatin, consistent with TF-mediated nucleosome remodeling [34].

Collectively, these results demonstrate that HiddenFoot can accurately infer TF footprints and nucleosome positioning from long-read Fiber-seq data, complementing SMF-based analyses. More importantly, HiddenFoot provides a principled approach to studying widespread heterogeneity in nucleosome organization across molecules and genomic regions—heterogeneity that may be driven or stabilized by TF binding. By enabling direct interrogation of nucleosome architecture at the single-molecule level, HiddenFoot offers a powerful framework for exploring the interplay between TF binding and chromatin dynamics in complex regulatory landscapes.

### Scope and Limitations of HiddenFoot

While HiddenFoot provides a powerful and flexible framework for interpreting single-molecule chromatin accessibility data, several modeling assumptions and biological factors present opportunities for further refinement and extension. First, the model assumes that TFs have residence times long enough to leave detectable footprints in the methylation patterns captured by SMF or Fiber-seq. However, labeling reactions in these assays typically last several minutes, whereas many TFs exhibit dwell times on the order of seconds. As a result, transient or fast-exchanging factors may be systematically underrepresented in the inferred binding profiles. Nonetheless, the local accessibility context near binding sites may provide indirect evidence of such binding events. For example, the absence of a nucleosome footprint covering a motif, combined with nucleosome footprints in flanking regions, can indicate TF occupancy. Future experiments that vary methyltransferase treatment times or combine SMF with live-cell single-molecule tracking (SMT) could help calibrate the sensitivity to residence time and expand the dynamic range of detectable footprints. Second, HiddenFoot models protein-DNA binding under thermodynamic equilibrium, ignoring the nonequilibrium effects of ATP-dependent processes such as chromatin remodeling, histone exchange, and transcription elongation. While this is a simplification, it is important to emphasize that the equilibrium model is used as a prior distribution over binding configurations. The actual methylation patterns, which serve as the observed data, strongly constrain the posterior distribution. Consequently, the inferred binding profiles can still reflect nonequilibrium features—such as cooperative co-binding or nucleosome eviction—even if these are not explicitly encoded in the energy model. This is analogous to our findings on TF-TF interaction energies, where patterns of cooperativity were uncovered despite not being explicitly modeled in the prior. Third, HiddenFoot requires PWMs to model sequence-specific binding preferences, limiting its applicability to TFs with known and well-characterized motifs. As a result, binding events from orphan or poorly characterized factors may go undetected. In future iterations, HiddenFoot could be extended to perform de novo motif discovery directly from binding footprints, leveraging the large number of inferred bound molecules and their sequence context. Such a capability would make the framework more general, enabling it to discover novel sequence-specific regulators or unanticipated binding modes.

Forth, HiddenFoot assumes that local protein concentrations are constant across all molecules mapping to the same genomic region. This simplifies the inference process and allows region-specific estimation of protein concentrations. However, in heterogeneous cell populations, molecules originating from cells in different cellular states may reflect varying concentrations of TFs or chromatin-associated proteins. In such cases, a more accurate model would account for a distribution of local concentrations within a single region. While this would increase biological realism, it also significantly raises model complexity and may introduce challenges related to overfitting and parameter identifiability. Addressing this limitation would likely require additional sources of information. A promising future direction could involve integrating SMF-like data with single-cell chromatin accessibility datasets, such as scATAC-seq, to better resolve cell-state–specific binding dynamics and concentration heterogeneity at regulatory loci.

By addressing these limitations—through experimental calibration, incorporation of nonequilibrium dynamics, motif-agnostic inference, and integration of multimodal data—HiddenFoot has the potential to evolve into a fully general platform for interpreting chromatin organization and transcriptional regulation at single-molecule resolution.

## Discussion

The advent of SMF and long-read chromatin profiling technologies such as Fiber-seq has fundamentally expanded our ability to study chromatin organization and regulatory logic at the resolution of individual DNA molecules. Yet the full potential of these technologies remains limited by the lack of computational tools capable of disentangling the overlapping and cooperative binding of TFs, nucleosomes, and Pol II from sparse and noisy methylation data. In this study, we introduce HiddenFoot, a biophysically grounded computational framework that overcomes these challenges by modeling protein-DNA binding as a thermodynamic process, enabling mechanistic interpretation of SMF and Fiber-seq data at single-molecule resolution.

HiddenFoot fills a critical gap in the current analytical landscape. Traditional approaches such as ChIP-seq, MNase-seq, or ATAC-seq provide population-averaged views of TF and nucleosome occupancy, masking the heterogeneity of chromatin states that exists across molecules and cells. More recent tools designed for SMF, including unsupervised methods like FootprintCharter [17], can stratify molecules by methylation patterns and detect footprints, but lack an explicit model of competitive binding and cannot resolve the molecular logic behind observed patterns. HiddenFoot advances the field by combining DNA sequence specificity, local concentration estimates, and methylation likelihoods within a statistical mechanics framework, producing molecule-wise binding probability maps for TFs, nucleosomes, and Pol II.

We demonstrate that HiddenFoot accurately infers TF footprints across thousands of loci, recapitulating known occupancy profiles for factors like CTCF and REST, while also identifying distinct and interpretable footprints for TFs with weak or diffuse ChIP-seq enrichment such as NRF1 and ESRRB. These results emphasize HiddenFoot’s ability to classify molecules into bound and unbound states, improving signal resolution over standard methylation-averaged footprints. The model is able to recover localized Pol II footprints in promoter regions, offering a complementary approach to recent Fiber-seq-based strategies such as FiberHMM [14], and provides a unified model for integrating Pol II, TFs, and nucleosomes on the same DNA molecule. A key contribution of HiddenFoot is its ability to systematically dissect TF-TF co-dependency. By estimating interaction energies between TF pairs from probabilistic single-molecule binding profiles, HiddenFoot distinguishes between co-occupancy driven by chromatin accessibility (e.g., nucleosome eviction) and direct physical cooperativity. Unlike correlation-based co-binding metrics, our approach is grounded in statistical thermodynamics and leverages null expectations derived from simulated equilibrium data. This allows us to identify TF pairs whose cooperative interactions are not explainable by chromatin-mediated effects alone, a necessary step for inferring direct regulatory interactions in complex gene networks. These findings complement previous work such as Sonmezer et al. [13], while extending interpretability by incorporating energy-based inference.

Beyond TFs, HiddenFoot reveals pervasive heterogeneity in nucleosome positioning, particularly evident in long-read Fiber-seq datasets from *Drosophila* S2 cells. By mapping nucleosome occupancy at the single-molecule level, we observe region-specific and molecule-specific variation in phasing and occupancy patterns around TF motifs. These results align with findings from Stergachis et al. [11], who documented widespread structural variability and nucleosome distortion linked to TF binding. Our analysis extends these insights by providing a probabilistic and quantitative measure of occupancy, enabling comparisons between TF-bound and unbound molecules. Importantly, we show that TF binding events such as MAX occupancy are associated with more ordered nucleosome phasing, consistent with the well-established role of MAX in promoteing chromatin remodeling [34].

A major strength of HiddenFoot lies in its flexibility and generalizability. The model performs robustly across distinct experimental platforms (SMF and Fiber-seq), organisms (mouse, human and *Drosophila*), and protein classes (TFs, nucleosomes and Pol II), using only input sequence, motif annotations, and binary methylation data. Its design is modular: additional biochemical constraints such as histone modifications, transcriptional output, or remodeling kinetics can be incorporated in future extensions. As new single-molecule and multi-omic platforms emerge, HiddenFoot offers a powerful, extensible foundation for interpreting their output in a mechanistic, interpretable framework.

Looking forward, HiddenFoot provides a versatile framework for dissecting cis-regulatory logic at single-molecule resolution. By resolving the combinatorial occupancy of TFs, nucleosomes, and Pol II on individual DNA molecules, it offers a powerful tool for investigating how regulatory elements function in different chromatin contexts. This capability is particularly valuable in systems where heterogeneity and cooperativity play central roles—such as during development, in disease progression, or in cell fate reprogramming—enabling deeper insight into how chromatin architecture shapes gene regulation across diverse biological conditions.

## Methods

### Biophysical Modelling of TF, Nucleosome, and Pol II Binding

HiddenFoot models TF, nucleosome, and Pol II binding as a competitive, probabilistic process governed by thermodynamic principles at equlibirum. Each DNA molecule is represented as a one-dimensional lattice, where protein-DNA binding configurations are evaluated using a Boltzmann distribution that incorporate both sequence specificity and local protein concentrations. Observed methylation patterns are linked to DNA accessibility through a probabilistic model that accounts for the stochastic nature of exogenous methyl-transferase activity.

### Binding energies and the Boltzmann factor

Each protein factor *n* is modeled by a sequence-specific binding energy function *E*_*n*_, which determines its affinity for a given DNA segment. For TFs, binding energies are computed from PWMs as:

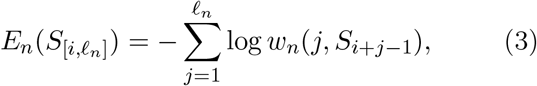

where *S*[*i,ℓ*_*n*_] denotes the DNA subsequence of length *ℓ*_*n*_ starting at position *i*, and *w*_*n*_(*j, b*) is the PWM entry for base *b* at position *j* for factor *n*. PWMs were obtained from the TF-clustered JASPAR2024 CORE database [20], which groups similar motifs to reduce redundancy.

For nucleosomes and RNA Polymerase II, we assumed no sequence specificity and modeled their binding as occurring over fixed-length segments of 147 bp and 40 bp, respectively. While weak sequence preferences for nucleosome positioning are known [35], we omitted these for simplicity in the present study.

Given a sequence segment *S*[*i,ℓ*_*n*_], the probability that factor *n* binds at position *i* is proportional to a Boltz-mann factor:

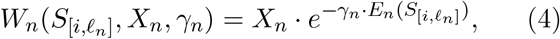

where *X*_*n*_ is an effective concentration parameter for factor *n*, and *γ*_*n*_ is a factor-specific inverse temperature parameter that modulates sequence selectivity. Higher values of *γ*_*n*_ yield steeper energy landscapes, favoring high-affinity sites, while lower values allow broader tolerance to sequence mismatches. For simplicity in this study, we assumed a common scaling factor (*γ*_*n*_ = *β*) across all proteins.

#### Binding configuration probability

Given a binding configuration *C*, which specifies a set of non-overlapping protein factors bound to subsequences of *S*, the probability of *C* under the model is proportional to the product of Boltzmann factors for all bound proteins in the configuration:

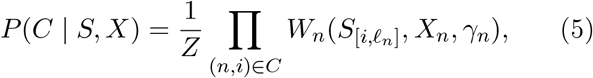

where (*n, i*) ∈ *C* denotes that protein factor *n* is bound at position *i* in configuration *C*, and *Z* is the partition function that normalizes over all allowed non-overlapping configurations.

Substituting the definition of the Boltzmann factors (Eq. 4) into the expression above, the configuration probability can be rewritten in terms of a total energy, recovering the standard Boltzmann-Gibbs form used in the main paper:

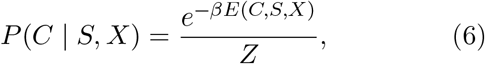

where the total energy is defined as:

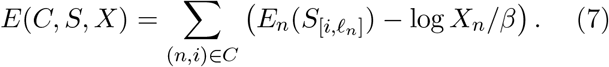

This expression shows that the probability of a configuration is determined by the sum of the binding energies of all bound proteins, with the logarithm of the concentrations acting as effective chemical potentials. This formulation ensures that high-probability configurations correspond to sets of strongly bound, non-overlapping factors, consistent with thermodynamic equilibrium.

### Partition function and dynamic programming

To efficiently evaluate the partition function *Z*, which sums over all possible non-overlapping configurations of bound proteins, we define a recursive partition function *Z*_*i*_ representing the total statistical weight of all valid configurations ending at position *i*:

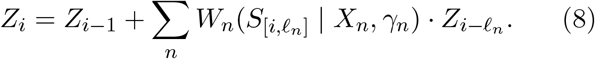

This recursion enables tractable computation over thousands of potential binding elements per region using dynamic programming.

### Incorporating methylation evidence

To relate binding configurations to observed single-molecule methylation patterns *M*_[*i,ℓ*]_, we introduce two likelihood models depending on whether the DNA segment is protected (bound) or unprotected (accessible):

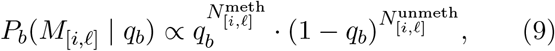

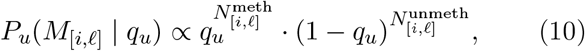

where *q*_*b*_ and *q*_*u*_ are the methylation probabilities for protected and unprotected DNA, respectively, and 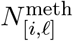 and 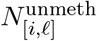 are the counts of methylated and unmethylated cytosines (or adenosines) in the segment. These statistics are calculated directly from the methylation data.

### Forward algorithm for marginal likelihood

To compute the likelihood of observing a methylation pattern *M* given a DNA sequence *S* and binding parameters, we combine the binding model with methylation likelihoods in a modified forward recursion:

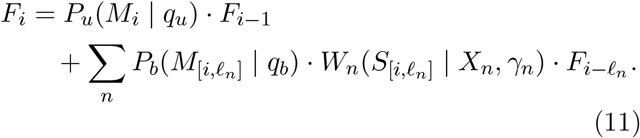

This forward pass computes the total joint likelihood of sequence-specific binding and methylation over the DNA molecule. The marginal log likelihood of the data is obtained as:

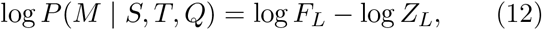

where *F*_*L*_ and *Z*_*L*_ are the forward likelihood and partition sum at the end of the molecule, respectively.

### Model Optimization and Binding Profile Inference

Parameter inference in HiddenFoot is achieved through two complementary approaches: maximum likelihood estimation using SGD, and posterior sampling using MCMC.

### SGD via Adam optimizer

The model maximizes the marginal likelihood of the observed methylation data across all molecules, given the DNA sequence and biophysical parameters. The objective is to maximize the log-likelihood (Eq. 12) with respect to the model parameters *θ* ∈ {log *X*_*n*_, *γ*_*n*_, *q*_*u*_, *q*_*b*_}. Gradients of the loglikelihood with respect to each parameter *θ*_*p*_ are computed analytically via dynamic programming:

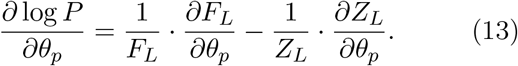

The derivatives of the forward recursion for *F*_*i*_ are given by:

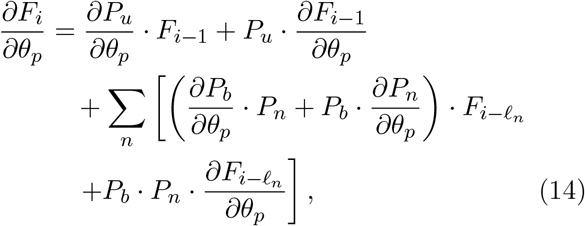

and for the partition function *Z*_*i*_:

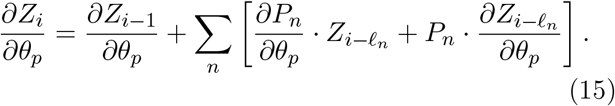

These recursions are evaluated iteratively for each position and in parallel across molecules using OpenMP. Gradients are accumulated in mini-batches and passed to the Adam optimizer [36] for stochastic gradient descent.

All parameters are internally transformed (e.g., using log or logit mappings) to constrain them to valid numerical domains and improve optimization stability.

### MCMC Sampling

To quantify parameter uncertainty, HiddenFoot optionally supports Bayesian inference via Metropolis–Hastings MCMC sampling [37]. Parameters are proposed using Gaussian distributions in transformed space, and accepted based on the Metropolis–Hastings ratio computed from the likelihood. This posterior sampling enables the computation of confidence intervals and model uncertainty.

### Binding probability computation

Once the model parameters are estimated, HiddenFoot computes the posterior binding probability for each factor at every candidate site using a forward–backward dynamic programming scheme. The forward algorithm computes the total statistical weight *F*_*i*_ of all configurations ending at position *i*, while the backward algorithm computes the complementary weight *B*_*i*_ of all configurations starting from position *i* + 1 to the end of the molecule.

For a given protein factor *n* and position *i*, the marginal posterior probability that *n* is bound at *i* on a particular molecule is given by:

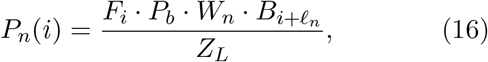

This formulation allows the model to marginalize over all compatible configurations, yielding accurate site-wise posterior probabilities that reflect both the DNA sequence context and the observed methylation pattern. For each factor, these per-molecule binding profiles *P*_*n*_(*i*) can be averaged across molecules to construct population-level footprints, or analyzed at single-molecule resolution to identify heterogeneous binding modes and cooperative interactions.

### Simulation framework

HiddenFoot includes a built-in simulation mode for generating synthetic single-molecule binding profiles and methylation datasets. Given an input DNA sequence and biophyscial paramters, the model first samples binding configurations from the Boltzmann-Gibbs equilibrium distribution. Configurations are sampled without overlap, respecting protein footprint lengths and steric exclusion constraints.

Given a sampled configuration, methylation states are then simulated using the probabilistic emission model: each cytosine (or adenosine, for Fiber-seq) is independently methylated or unmethylated depending on whether it is protected by a bound protein, with probabilities determined by the parameters *q*_*U*_ and *q*_*B*_. This mimics the exogenous methyltransferase labeling process used in SMF and Fiber-seq experiments.

Simulations can be run using user-defined protein concentrations and energy parameters, allowing precise control over occupancy dynamics in synthetic data. Simulated datasets can be generated at scale for arbitrary sequences, motif sets, and fitted or hypothetical parameter values.

Once model parameters are learned from real data, HiddenFoot’s generative mode can be used to simulate binding and methylation profiles under thermodynamic equilibrium, independent of the observed methylation signal. This provides a powerful way to compare predicted equilibrium behavior to experimental data, enabling the identification of deviations from equilibrium due to biological energy expenditure (e.g., chromatin remodeling or transcription), or cooperative interactions not explicitly modeled in the energy function.

### Differentiating Physical and Nucleosome-Mediated TF Cooperativity

#### Inference of TF-TF interaction energies

To determine whether TF pairs exhibit cooperative, repulsive, or independent binding behavior, we infer a shared binding energy term *E*_12_ based on single-molecule cobinding patterns. This interaction term quantifies how the binding of one TF influences the binding of another on the same DNA molecule.

We define four occupancy classes for a given TF pair across all DNA molecules: *N*_00_, *N*_10_, *N*_01_, and *N*_11_, representing the number of molecules in which TF1 and TF2 are unbound or bound, respectively. Here, a subscript digit of 1 indicates binding and 0 indicates the absence of binding for each TF, so that, for example, *N*_10_ corresponds to molecules where TF1 is bound and TF2 is unbound.

Unlike discrete classification, we compute these values using the marginal binding probabilities inferred by HiddenFoot. For each molecule *m*, we use the posterior binding probabilities 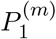 and 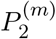 of TF1 and TF2, respectively, and approximate the expected cooccupancy counts across all molecules as:

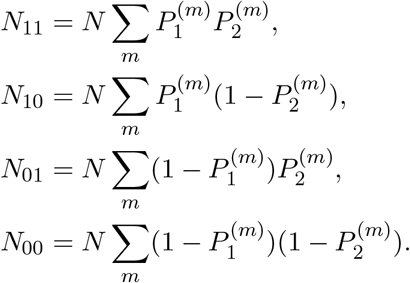

where *N* is the total number of molecules.

These quantities represent the expected number of molecules in each co-binding state, marginalized over probabilistic assignments rather than hard labels.

We model the probability of each co-binding state based on the individual TF binding energies *E*_1_, *E*_2_, and their shared interaction energy *E*_12_:

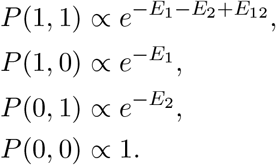

The partition function normalizing these probabilities is given by:

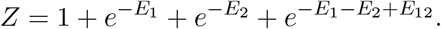

The overall log-likelihood of observing the expected co-binding pattern is:

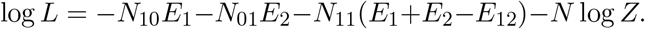

To estimate the optimal energy parameters, we minimize the negative log-likelihood using the Nelder–Mead simplex optimization algorithm. The key quantity of interest is *E*_12_:

- *E*_12_ ≈ 0: implies independent binding,
- *E*_12_ *<* 0: indicates mutually exclusive binding,
- *E*_12_ *>* 0: indicates cooperative co-binding.

To improve convergence and avoid local minima, we initialize the optimization using optimal binding energies assuming no interaction: 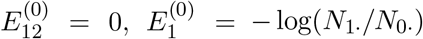 and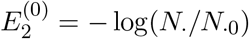.

The initialized values are then refined by minimizing the full log-likelihood function with respect to *E*_1_, *E*_2_, and *E*_12_. The resulting shared energy term *E*_12_ provides a robust, quantitative measure of TF-TF interaction captured from probabilistic single-molecule binding data.

#### Comparing interactions energies from real and simulated binding profiles

To distinguish direct physical TF-TF interactions from co-binding effects mediated by nucleosome organization or steric hindrance, we used simulations generated with HiddenFoot’s equilibrium generative model. These simulations apply the same protein-specific binding parameters inferred from real data, but exclude any explicit TF-TF interaction terms.

In this setting, any observed co-dependency arises solely from steric exclusion or nucleosome competition, as no energetic coupling between TFs is encoded. By repeating the interaction energy inference procedure described above on both real and simulated datasets, we can identify TF pairs whose co-dependencies are consistent with structural effects versus those indicative of biochemical interactions.

TF pairs that exhibit strong interaction energies in real data but not in the corresponding simulated datasets are interpreted as having true physical cooperativity. In contrast, pairs with similar shared energies across both conditions likely reflect chromatin-mediated co-binding constraints.

All comparisons were made using matched genomic loci and motif positions, ensuring consistency across real and simulated contexts.

### Implementation and code avilability

HiddenFoot is implemented in C++ and uses OpenMP for multithreading across molecules. The code includes both forward and backward recursions for computing the partition function and posterior probabilities, with analytical gradient support for parameter optimization via SGD using the Adam optimizer, and MCMC for estimating parameter uncertainties. Its modular design supports multiple execution modes (e.g., training, inference, simulation), controlled via command-line options, enabling fast analysis of thousands of molecules and motifs per genomic region. The code is freely available on GitHub: https://github.com/MolinaLab-IGBMC/HiddenFoot

### SMF and Fiber-seq Datasets and Methylation Calling

#### Methylation calling of *Drosophila* S2 cells

We retrieved methylation data for *Drosophila* S2 cells from the GSE146942 dataset [11]. Fiber-seq data was processed to extract methylation information using the M6A methyltransferase signal on adenine/thymine bases. Custom R scripts were used to parse PacBio reads and convert them into binary methylation matrices at single-base resolution.

#### Methylation calling of mESCs

BAM files for mESC single-molecule footprinting were obtained from Sonmezer et al. [13]. A custom R script was developed to extract the methylation state of each cytosine within accessible regions. The data was binarized and formatted into molecule-wise matrices representing methylated and unmethylated CpG/GpC sites.

#### Extracting Methylation information for regions of interest

To investigate specific TF footprints, we identified genomic regions containing relevant motifs using FIMO. Accessible regions were obtained from ATAC-seq data (*Drosophila*) and DNAse-seq data (mESC). We extended +/-500 bp windows (*Drosophila*) or +/-400 bp (mESC) around each motif instance. A custom R pipeline extracted methylation data and sequence content for each region, assembling matrices for use with HiddenFoot.

### Validation of HiddenFoot TF Binding Predictions Using Bulk Experiments

To validate TF binding predictions, we compared HiddenFoot-inferred binding probabilities with ChIP-seq signal intensity across the genome. For each region, we binned TF motifs by ChIP-seq peak height and calculated average binding probabilities within each bin. Pearson correlation coefficients between HiddenFoot binding scores and log-transformed ChIP-seq signal were computed per TF. This approach was used for CTCF, REST, NRF1, and ESRRB in mESCs.

We compared HiddenFoot-predicted nucleosome occupancy profiles to MNase-seq signal in both S2 cells and mESCs. MNase-seq data in BigWig format was processed to extract the maximum signal within +/-500 bp around each region of interest. We computed the average nucleosome probability predicted by HiddenFoot for each region, binned regions by MNase signal, and calculated per-bin means. Pearson correlation coefficients between binned MNase and model predictions were used to assess agreement.

### Public datasets used for validation and analysis

| Study | Cell Type | Data Type |
| --- | --- | --- |
| GSE172392 | mESC | MNase-seq |
| GSE169222 | S2 | MNase-seq |
| GSE96581 | S2 | CTCF ChIP-seq |
| GSE46614 | S2 | CLAMP ChIP-seq |
| GSE55306 | S2 | MAX ChIP-seq |
| GSE30206 | mESC | CTCF ChIP-seq |
| GSE27148 | mESC | REST ChIP-seq |
| GSE11431 | mESC | ESRRB ChIP-seq |
| GSE67867 | mESC | NRF1 ChIP-seq |
| GSE274357 | mESC | ATAC-seq |
| GSE103177 | *Drosophila* S2 | ATAC-seq |

## Acknowledgments

This work was carried out at the Interdisciplinary Thematic Institute IMCBio+, part of the ITI 2021–2028 program of the University of Strasbourg, CNRS, and Inserm. It was supported by IdEx Unistra (ANR-10-IDEX-0002), the SFRI-STRAT’US project (ANR-20-SFRI-0012), and EUR IMCBio (ANR-17-EURE-0023), under the framework of the France 2030 Program. LD was supported by the IMCBio PhD program, and HM was supported by the Région Grand Est through the program Projects 2021 / Action 15: Volet 2 Compétences Recherche: Doctorants et Jeunes Chercheurs, as well as by the Agence Nationale de la Recherche (ANR) project ANR-20-CE12-0014. This work was conceived during NM’s sabbatical stay in the Krebs lab, made possible through the support of the Theory@EMBL program. NM is especially grateful to Arnaud Krebs for his hospitality and the stimulating discussions. Language editing and proofreading assistance was provided by ChatGPT (OpenAI).

## Supplementary Information

**Figure S1:**
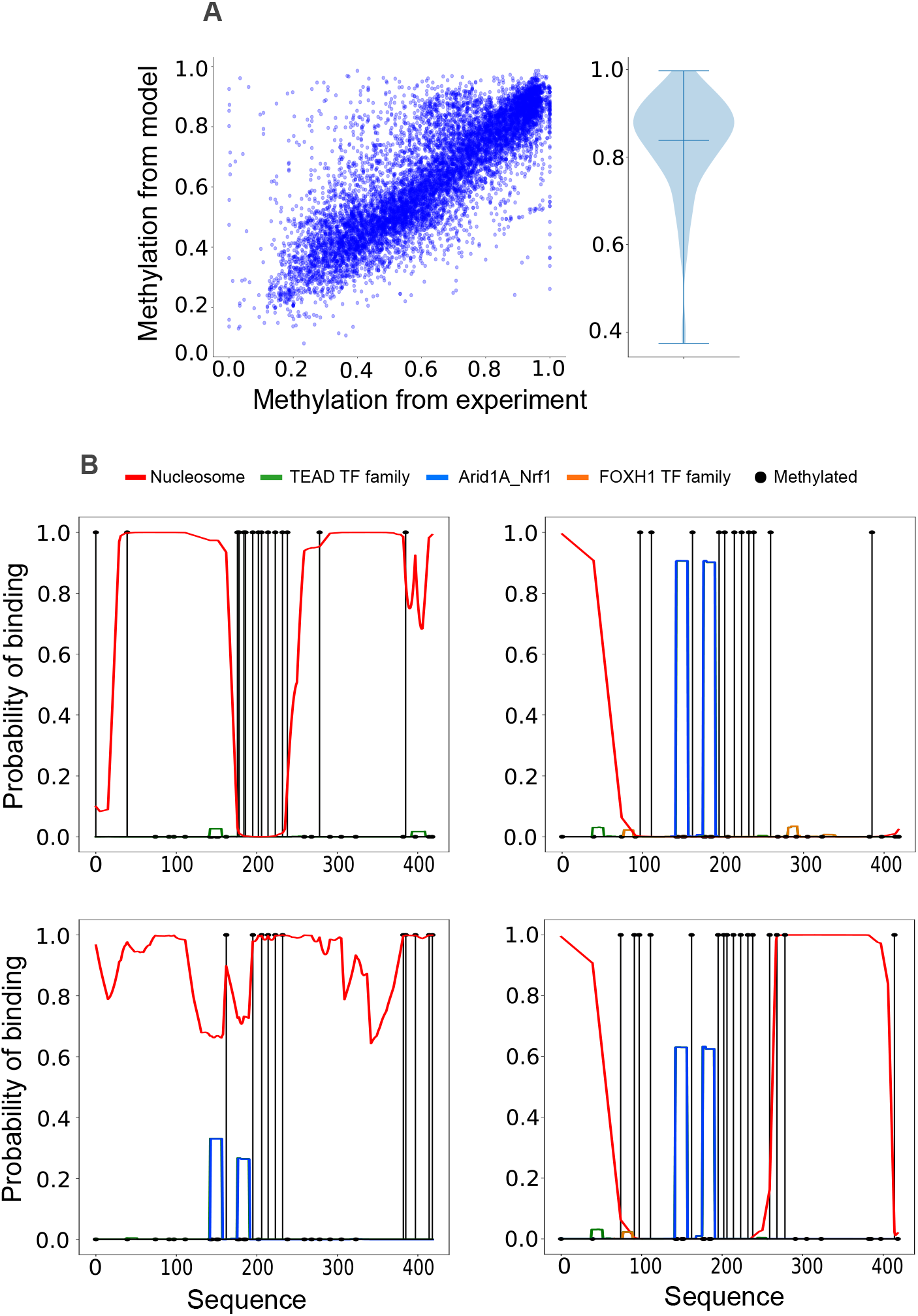
Model-predicted methylation and binding profiles across amplicon regions. **(A)** Correlation between predicted and observed methylation patterns. The left panel shows a scatter plot comparing the average methylation level at each cytosine site across all amplicon regions, as predicted by the HiddenFoot model versus experimentally observed methylation. Each point represents a single site. The right panel displays a violin plot summarizing the Pearson correlation coefficients between predicted and observed methylation across individual amplicons, indicating strong predictive performance. **(B)** Representative example of single-molecule predictions. Each row corresponds to an individual molecule from a high-coverage amplicon, with vertical stems indicating the methylation state at each cytosine (black for methylated, orange for unmethylated). Overlaid are the inferred posterior binding probabilities from HiddenFoot, with nucleosome occupancy shown in red and NRF1 TF binding shown in green. This visualization highlights the model’s capacity to accurately reconstruct both sparse methylation data and complex protein-DNA interaction patterns at single-molecule resolution.

**Figure S2:**
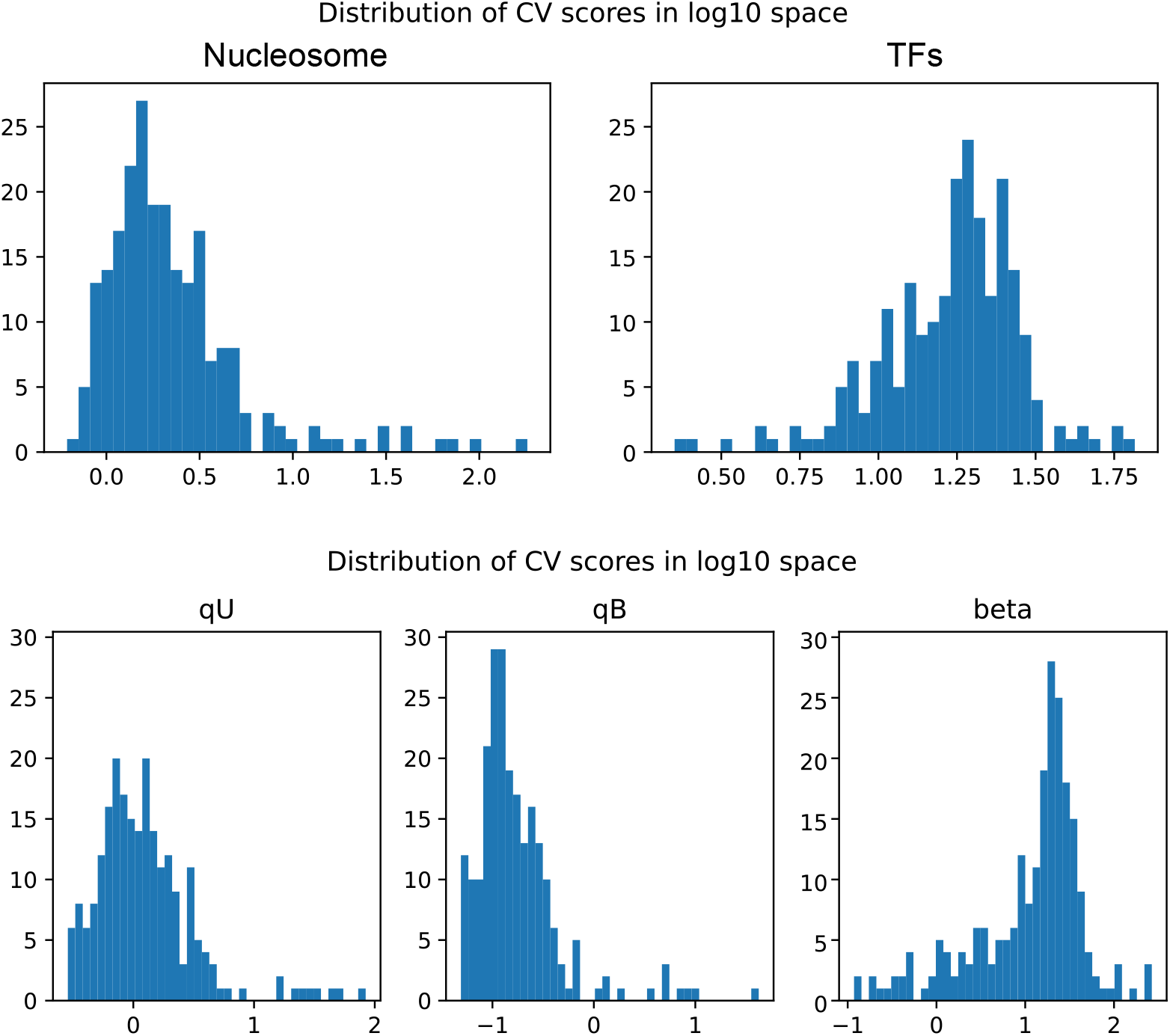
Posterior variability of model parameters across amplicon regions. Distributions of the coefficient of variation (CV), defined as the standard deviation divided by the mean, for parameters estimated via posterior sampling using Markov Chain Monte Carlo (MCMC). Each distribution summarizes results across all amplicon regions analyzed with HiddenFoot. Parameters include nucleosome and TF concentrations, as well as the inverse temperature parameter *β*. In general, CVs are low, indicating robust parameter identifiability. Nucleosome concentrations show particularly high precision, with CVs consistently below 10%. TF concentrations and *β* also exhibit good identifiability, with most CVs below approximately 30%. These results support the identifiability of the model’s parameters.

**Figure S3:**
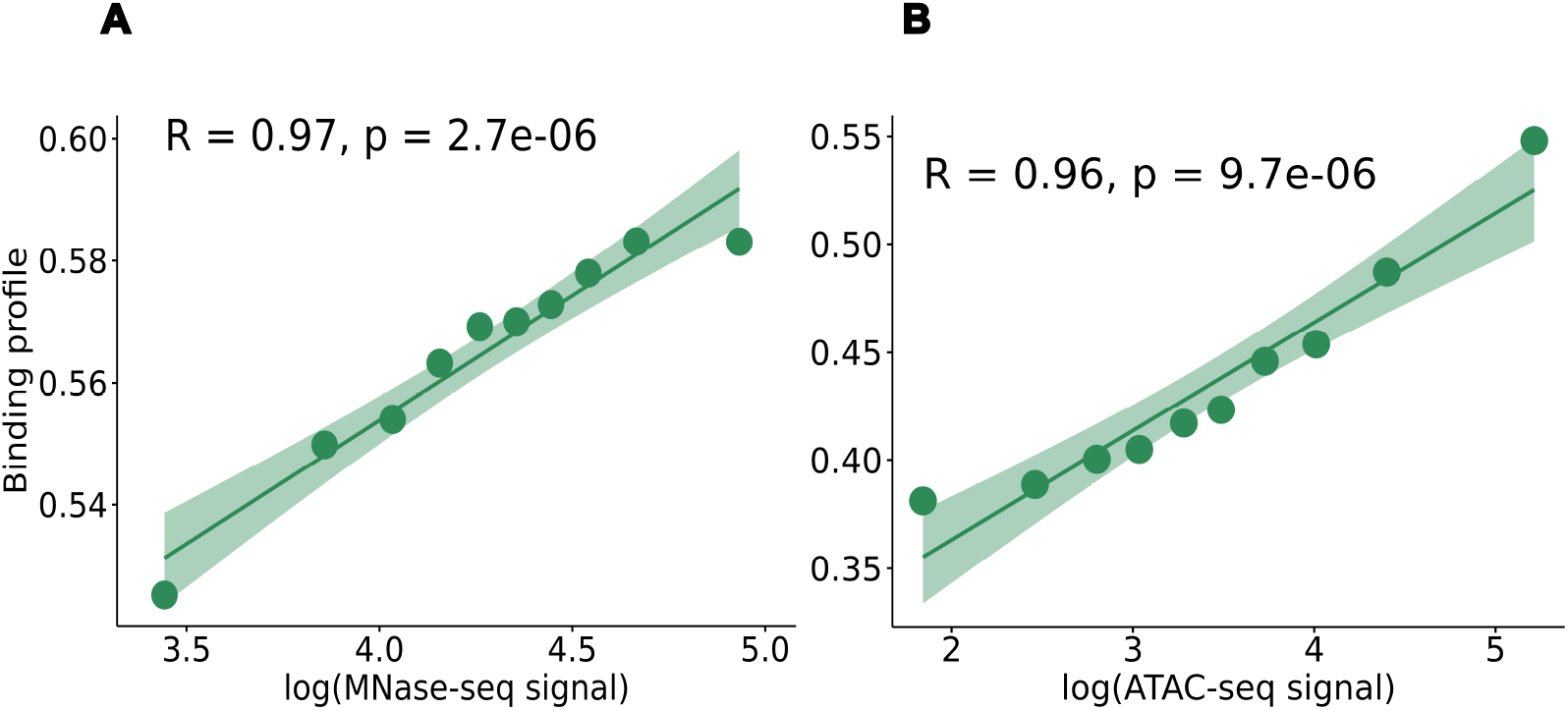
HiddenFoot-inferred accessibility correlates with independent chromatin accessibility measurements. **(A)** Correlation between HiddenFoot-predicted probability of DNA accessibility and MNase-seq signal (log-transformed). **(B)** Correlation between HiddenFoot-predicted accessibility and ATAC-seq signal (log-transformed). Each dot represents the average across genomic regions grouped into decile (10-quantile) bins, sorted according to increasing MNase-seq or ATAC-seq signal. Shaded areas indicate the 95% confidence interval of the linear fit. Pearson correlation coefficients (*R*) and associated *p*-values are reported in each panel, showing strong agreement between HiddenFoot predictions and experimental accessibility measures (*R* = 0.97, *p* = 2.7 × 10^*−*6^ for MNase-seq; *R* = 0.96, *p* = 9.7 × 10^*−*6^ for ATAC-seq).

**Figure S4:**
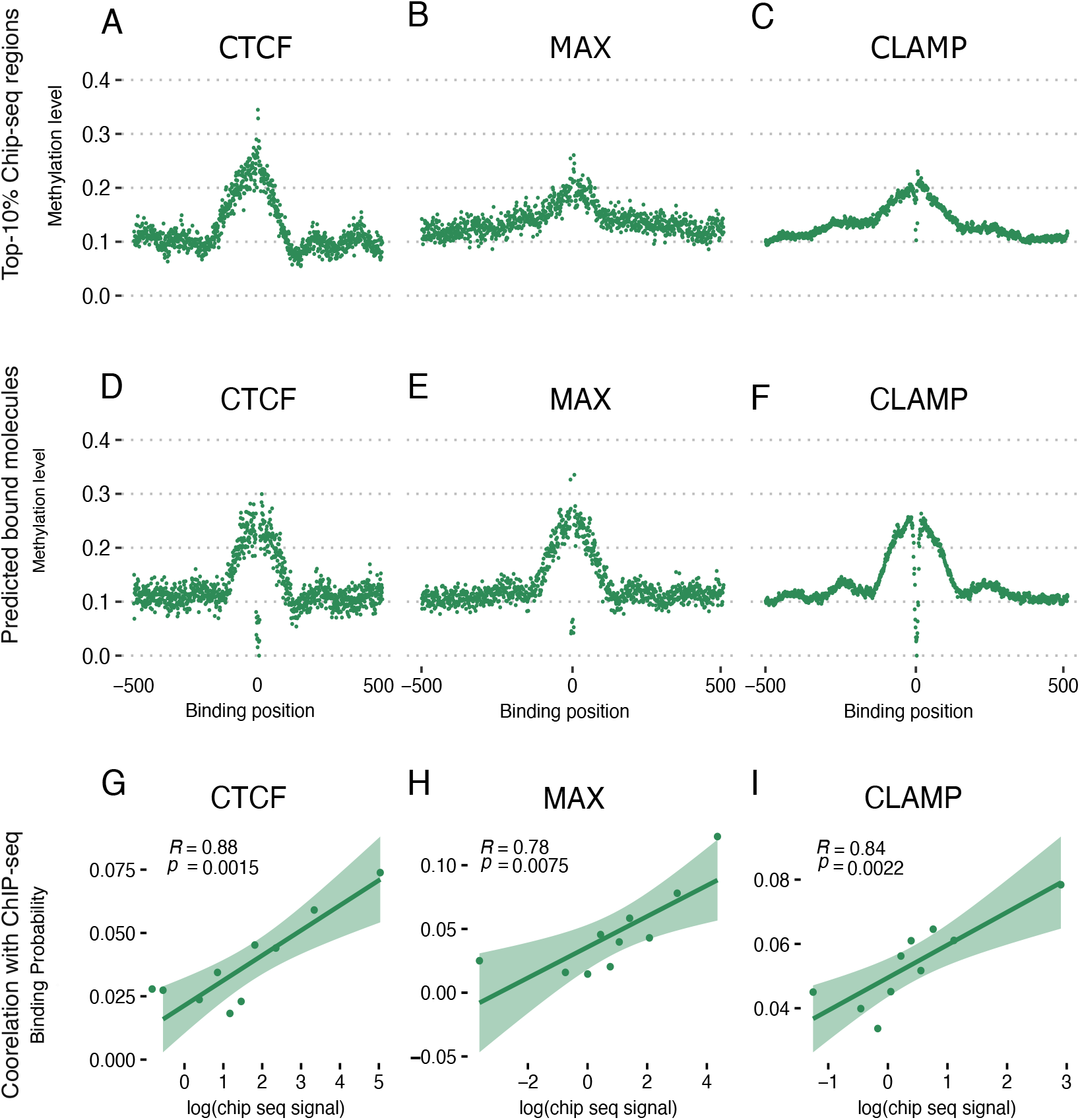
HiddenFoot detects methylation footprints in Drosophila S2 cells using Fiber-seq. **(A–C)**: Average methylation level at the binding site of the top 10% ChIP-seq enriched molecules for CTCF (A), MAX (B), and CLAMP (C). Each dot represents the average methylation level at a given position relative to the center of the TF motif. **(D–F)**: Average methylation levels at the same TF motifs, stratified using HiddenFoot predictions. Molecules are divided into bound and unbound categories based on inferred binding state. Clear footprint-like patterns are visible, particularly for CTCF and CLAMP. **(G–I)**: Pearson correlation between the average HiddenFoot-predicted binding probabilities and log-transformed ChIP-seq signal, binned into 10 groups. Each dot represents one bin. HiddenFoot predictions show strong correlation with ChIP-seq signal for CTCF (*R* = 0.88, *p* = 0.0015), MAX (*R* = 0.78, *p* = 0.0075), and CLAMP (*R* = 0.84, *p* = 0.0022), supporting the model’s accuracy in predicting TF occupancy from single-molecule methylation data in Drosophila.

**Figure S5:**
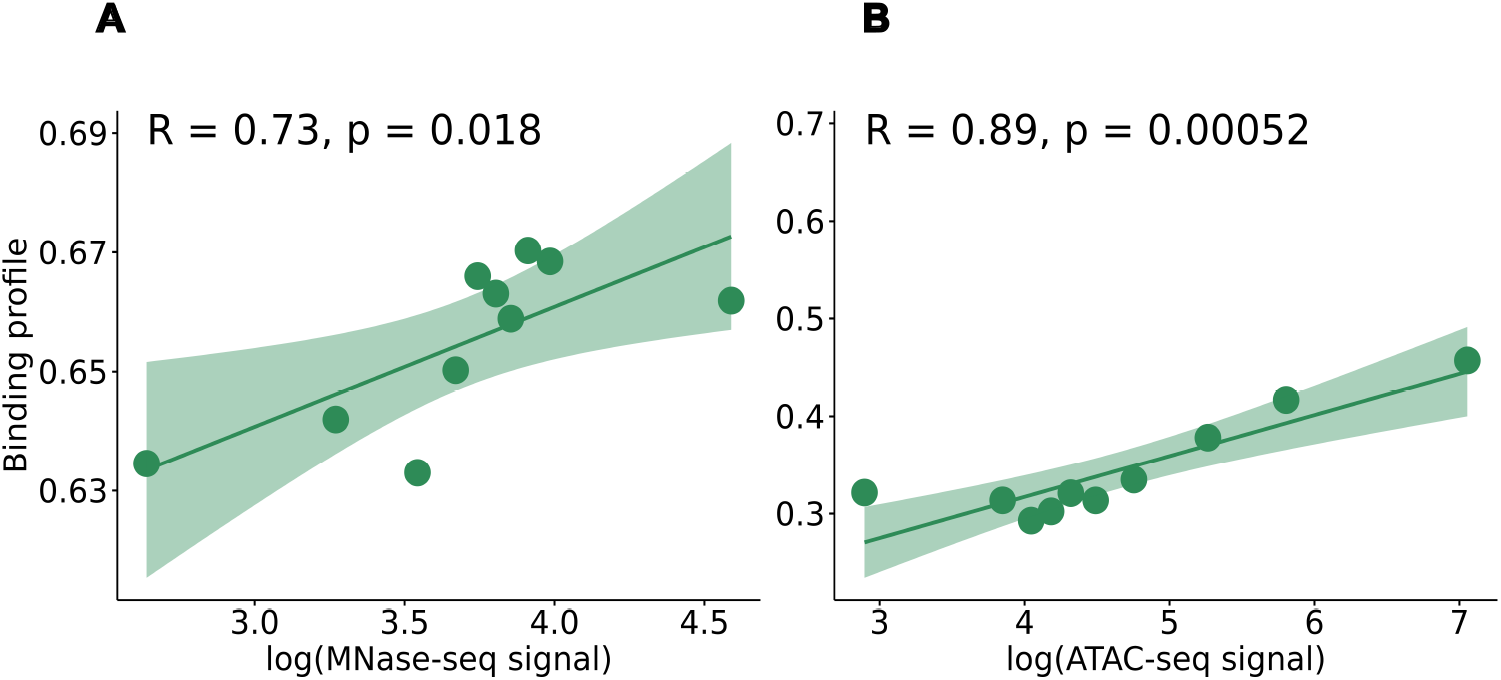
HiddenFoot-inferred accessibility correlates with chromatin accessibility measurements in S2 cells using Fiber-seq. **(A)** Correlation between HiddenFoot-predicted probability of DNA accessibility and MNase-seq signal (log-transformed).**(B)** Correlation between HiddenFoot-predicted accessibility and ATAC-seq signal (log-transformed). Each dot represents the average across genomic regions grouped into decile (10-quantile) bins, sorted according to increasing MNase-seq or ATAC-seq signal. Shaded areas indicate the 95% confidence interval of the linear fit. Pearson correlation coefficients (*R*) and associated *p*-values are reported in each panel, showing strong agreement between HiddenFoot predictions and experimental accessibility measures in S2 cells (*R* = 0.73, *p* = 0.018 for MNase-seq; *R* = 0.89, *p* = 5.2 × 10^*−*4^ for ATAC-seq).

**Figure S6:**
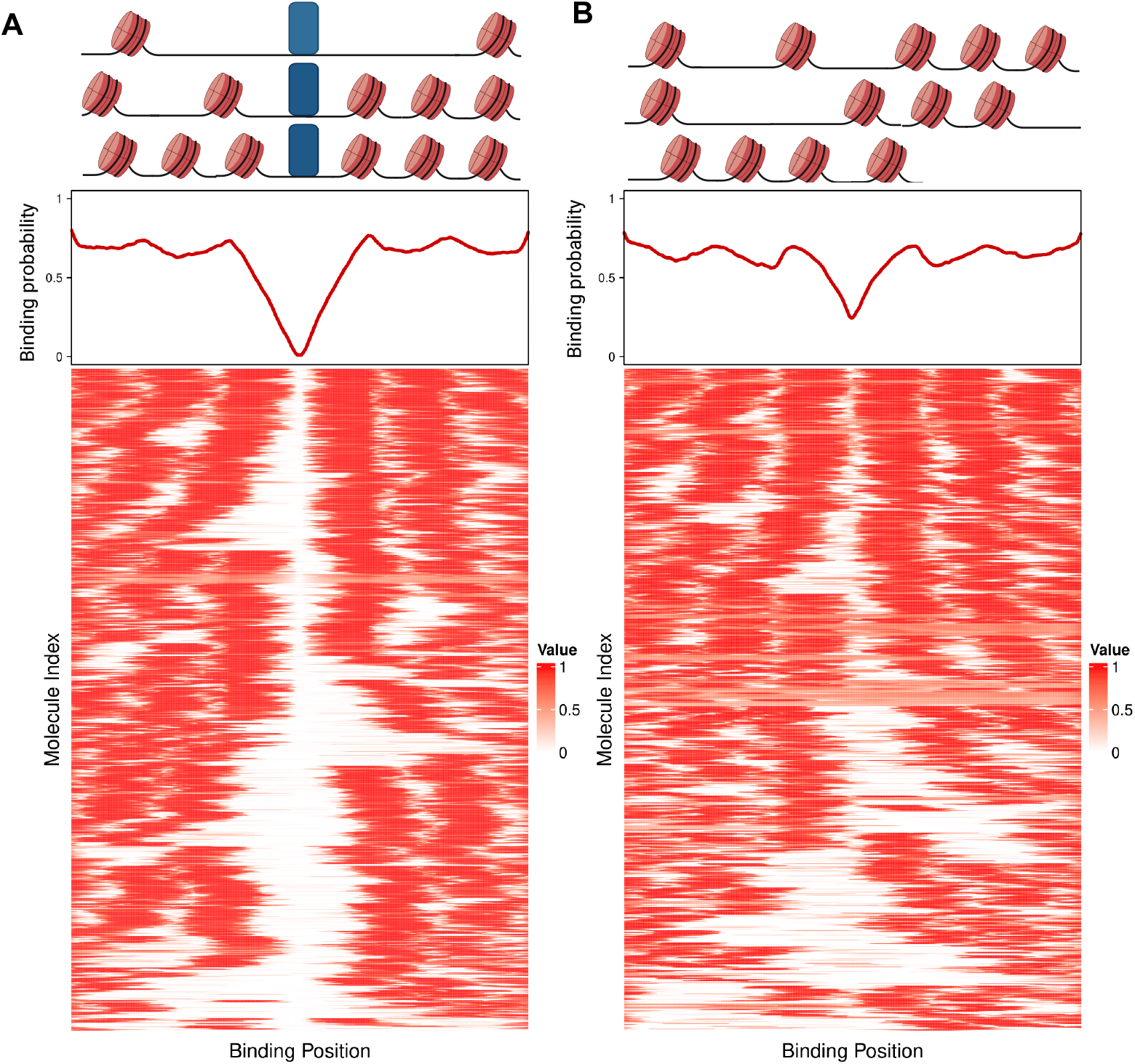
Figure S6. HiddenFoot reveals heterogeneous nucleosome positioning around MAX binding sites. Heatmaps display single-molecule occupancy profiles centered on MAX TF motifs across the genome, clustered by similarity in nucleosome positioning. **(A)** Molecules predicted to be bound by MAX. **(B)** Molecules predicted to be unbound. Each row represents an individual molecule, with colors indicating the posterior probability of nucleosome occupancy. Diagrams above the heatmaps illustrate the hypothesized influence of MAX binding on local nucleosome organization. In both cases, HiddenFoot reveals high heterogeneity in nucleosome structure and phasing, with distinct patterns emerging between MAX-bound and unbound subpopulations. These results suggest that MAX binding locally shapes chromatin architecture, while unbound regions retain more variable nucleosome configurations.

